# Keratinocyte Arginase1 Deficiency Impairs Healing and Antimicrobial Defence against *Pseudomonas aeruginosa* Infection

**DOI:** 10.1101/2025.02.24.640016

**Authors:** Rachel A. Crompton, Denis C. Szondi, Christopher Doherty, Helen A. Thomason, Sze Han Lee, Yan Ping Loh, Catherine A. O’Neill, Leah Vardy, Andrew J. McBain, Sheena M. Cruickshank

## Abstract

Wound infection is a major disruptor of wound healing. Keratinocytes, critical in repair and microbial responses, require the L-arginine hydrolysing enzyme arginase1, for effective healing. Wound pathogens such as *Pseudomonas aeruginosa* may also need L-arginine. We therefore investigated host-microbial interactions in the context of wound healing and L-arginine metabolism. Arginase-inhibited murine wounds challenged with *P. aeruginosa,* exhibited significantly delayed re-epithelialisation. This finding was recapitulated *in vitro* using *P. aeruginosa-*challenged, arginase1 deficient (*shARG1)* keratinocytes, associated with reduced epithelial proliferation and viability, and heightened inflammation. Whilst *P. aeruginosa* challenge promoted host metabolism of L-arginine, this was perturbed in wounded *shARG1* keratinocytes. There was, however, heightened downstream polyamine metabolism in *shARG1* cells when under *P. aeruginosa* challenge. Host keratinocyte arginase1 deficiency promoted bacterial growth *in vitro*, in line with a failure to upregulate the anti-microbial peptides, *β-defensins,* in *shARG1* scratches. This work demonstrates a pivotal role for keratinocyte arginase1 in wound infection.

**HIGHLIGHTS:**

1. Host arginase is required for effective healing under *P. aeruginosa* challenge.
2. *P. aeruginosa* enhances host keratinocyte L-arginine metabolism upon scratch.
3. *P. aeruginosa* promotes polyamine metabolism in arginase1 deficient wounds *in vitro*.
4. Arginase1 is required for keratinocyte anti-microbial defence against *P. aeruginosa*.

## INTRODUCTION

Chronic wounds are a major debilitating disorder, often associated with advanced age and underlying metabolic disorders, such as diabetes ^1^. As we age, metabolic changes occur, including reduced cutaneous expression of the enzyme arginase 1 (ARG1) ^2^. This is important as ARG1 is upregulated throughout the healing process and is required for effective healing ^3–7^. Reduced ARG1 expression is associated with delayed healing and a ‘non-healing’ outcome of chronic foot ulcers ^3,8^.

In mammalian cells, ARG competes with several substrates to catabolise the amino acid L-arginine, resulting in the production of metabolites including; arginine glycine amidotransferase (AGAT) producing L-ornithine and creatine; nitric oxide synthase (NOS) that produces NO and L-citrulline; and ARG, which converts L-arginine to urea and L-ornithine. Metabolism of L-ornithine can subsequently generate L-proline, L-glutamate, polyamines and L-citrulline ^9,10,11,12^ (Figure S1a). ARG activity is associated with inflammation resolution, cell proliferation, differentiation and collagen synthesis ^2,8,13,14^. An important source of ARG1 in the skin are keratinocytes - the main cell type of the epidermis. These multi-functional cells are important immune regulating cells essential for skin barrier maintenance, orchestration of the healing response and production of anti-microbial products ^15–17^. Wounds are innately subject to contamination by environmental microbes and susceptible to infection ^18,19^. L-arginine metabolism contributes to wound defence via NO production ^20^ and via ARG1 mediated-metabolism of polyamines ^21,22^ or L-proline ^23^. However, L-arginine and its metabolic products can also benefit microbes, as certain bacteria can utilise these metabolites, in bacterial respiration, fermentation and replication ^24,25^. *Pseudomonas aeruginosa,* a bacterium commonly found within chronic wounds ^26^, can utilise L-arginine to produce energy in the form of ATP and promote its growth via putrescine synthesis ^24^.

Given ARG activity is necessary for effective healing and can also impact bacterial metabolism, we considered how infection and ARG deficiency, associated with human ageing, impact both the healing response and bacterial colonisation. This study aimed to determine the role of ARG1 in the cutaneous wound and keratinocyte response to *P. aeruginosa* challenge, using a combination of *in vitro* and *in vivo* model systems.

## RESULTS

### Inhibition of host ARG exacerbated the *P. aeruginosa* induced delay in cutaneous healing

mRNA levels increased in wounded keratinocytes, 6h post co-culture with *P. aeruginosa in vitro* (Figure S1b) suggesting a potential role of ARG1 in wound infection, Murine excisional wounds were subcutaneously injected with the ARG inhibitor nor-NOHA and / or topically applied *P. aeruginosa,* and collected 3 days post wounding (Figure 1a-b, Figure S1c). Histo-morphometric quantification revealed that in combination, a significant delay in re-epithelialisation was observed (Figure 1a, b). This effect was recapitulated in *in vitro* HaCaT keratinocyte scratches (Figure 1c) and *shARG1* knock down NTERT-1 keratinocytes (Figure 1d). These results demonstrate a lack of ARG1 is detrimental to the cutaneous healing response during wound infection.

**Figure 1.**
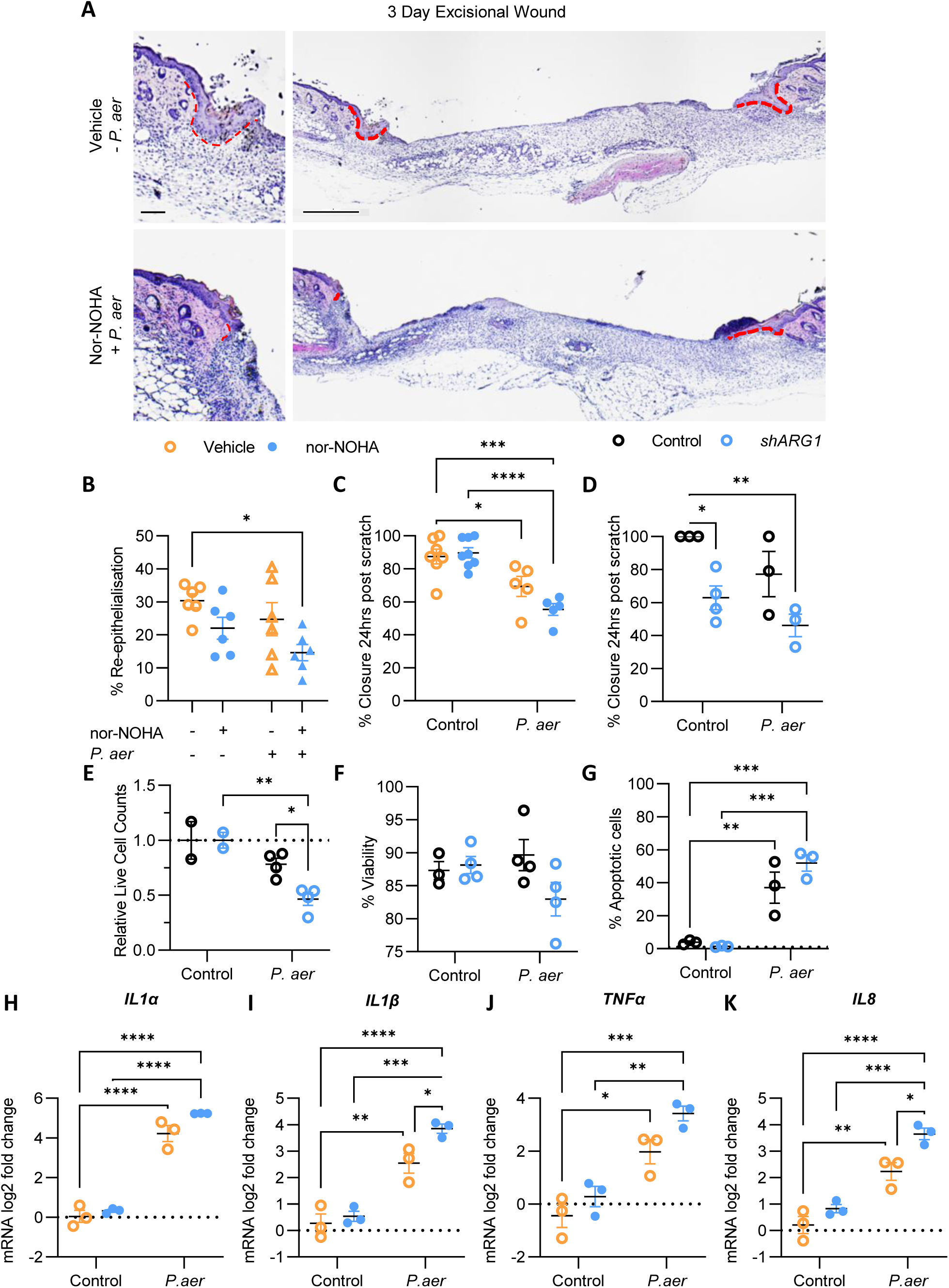
A *P. aeruginosa* induced delay in cutaneous healing is exacerbated by host inhibition of arginase, associated with reduced keratinocyte proliferation and heightened apoptosis and inflammation. (A) H&E-stained representative images illustrating reduced re-epithelialisation of day 3 excisional murine wounds topically treated with *P. aeruginosa* and subcutaneously injected with arginase inhibitor nor-NOHA compared to control. Red line indicates length of re-epithelialisation from the wound edge (full wound scale = 500µm, higher magnification image scale = 100µm), quantified in (B); n=6/group. (C) Quantification of HaCaT scratch wounds 24h post scratch and co-culture with *P. aeruginosa* and nor-NOHA, expressed as percent closure relative to initial scratch area; n=6-8/group. (D) Quantification of NTERT-1 sh*ARG1* knock down keratinocyte scratch closure co-cultured with *P. aeruginosa* (10^4^ cfu/ml) 24h post scratch; n=3-4/group. (E-F) NTERT-1 sh*ARG1* cell counts and viability 24h post co-culture with *P. aeruginosa*, determined by live/dead AO/PI staining; n=3-4/group. (G) Flow cytometric analysis of sh*ARG1* NTERT-1 keratinocytes stained with Annexin-V for apoptosis assessment; n=3/group. mRNA analysis by qPCR of cytokines *IL1α* (H), *IL1β* (I), *TNFα* (J) and chemokine *IL8* (K), in HaCaT cells scratched and co-cultured for 24hrs with *P. aeruginosa,* and/or ARG inhibited with nor-NOHA. N = 3/group. Data presented as log 2^-ΔΔCt relative to control. Data presented as individual points graphed with mean +/- SEM; p<0.05=*, p<0.01=**, p<0.001=***, p<0.0001=****. See also Figure S1.

Keratinocyte proliferation and viability are important for re-epithelialisation ^27^. Quantification of live/dead cell counts of *shARG1* keratinocytes, co-cultured with *P. aeruginosa* for 24hrs, showed significantly reduced live cell counts compared to either uninfected *shARG1* or control cells cultured with *P. aeruginosa* (Figure 1e). A similar trend was observed with reduced viability and increased apoptosis of *shARG1* cells co-cultured with *P. aeruginosa,* compared to controls, verified with flow cytometry (Figure 1f, g). *P. aeruginosa* also induced significantly increased *IL1α, IL1β, TNFα* and *IL8* mRNA levels in keratinocytes (Figure 1h-k), notably the highest cytokine expression was observed in the *P. aeruginosa* and nor-NOHA treated scratched keratinocytes, of which *IL1β* and *IL8* were most significant (Figure 1i, k).

### *P. aeruginosa* promotes host L-arginine metabolism in keratinocytes

To understand how the L-arginine pathway (Figure S1a) is altered in bacterially challenged ARG inhibited keratinocytes, key L-arginine metabolic enzymes and metabolite levels were assessed by qPCR and liquid chromatography–mass spectrometry (LC-MS).

Upon bacterial challenge intracellular keratinocyte L-arginine levels were significantly reduced, more so upon scratch (Figure 2a). Levels of the enzyme *NOS2* (Figure 2b) mRNA were higher in *P. aeruginosa-*challenged keratinocytes, and further increased on scratch. This corresponded with higher levels of the NOS downstream product L-citrulline (Figure 2c). However, despite upregulation of keratinocyte *ARG1* mRNA by *P. aeruginosa* upon scratch (Figure 2d), no difference was observed in the ARG products, L-ornithine (Figure 2e) and urea (Figure 2f). L-arginine can also be metabolised by arginine decarboxylase (ADC) to form agmatine ^28^; however, whilst the concept of a mammalian ADC has been established ^29,30^, the presence of a human ADC in skin and keratinocytes is unclear. We found that in a sterile environment, keratinocytes produce marginally higher levels of agmatine upon scratch; however, this was significantly higher upon *P. aeruginosa* challenge (Figure 2g). Agmatine generation from L-arginine via bacterial ADC is well-established ^31^. Indeed, whole genome sequencing of the *P. aeruginosa* NCTC10781 strain used in this study showed metabolic regulation of several pathways, comprising many L-arginine pathway-related genes (Figure S1 a, S2a-b, Table S1, 2), including the ADC gene *adiA/speA* (EC:4.1.1.19) (Table S1, 2). Collectively this suggests *P. aeruginosa* is promoting a higher degree of L-arginine metabolism in wounded keratinocytes predominantly via NOS and ADC activity.

**Figure 2.**
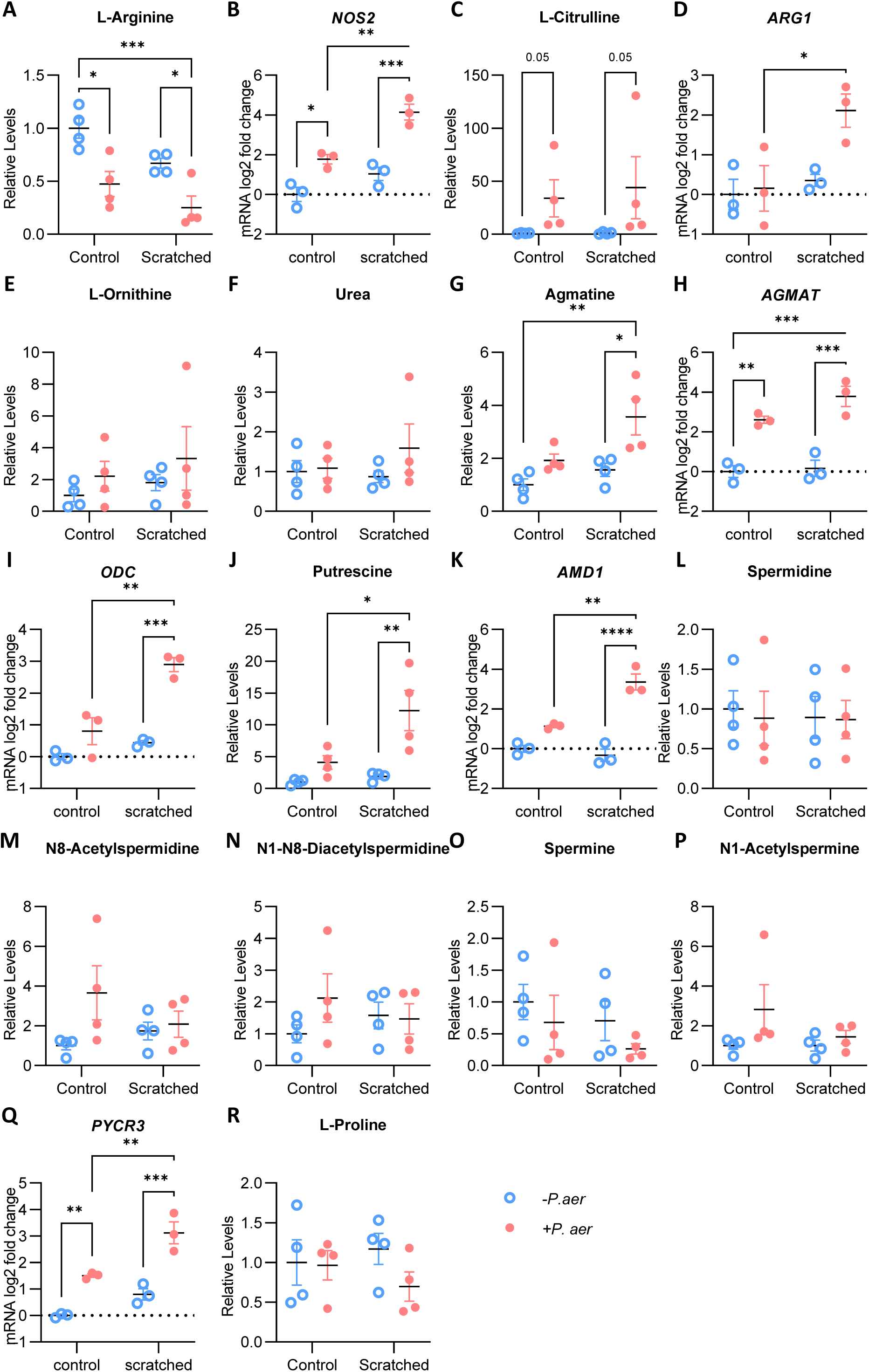
*P. aeruginosa* increases host arginine metabolism in keratinocytes upon wounding. Quantification of metabolite levels in unscratched v scratched keratinocytes, 12h post co-culture with *P. aeruginosa*, determined by LC-MS. Data presented as metabolite levels relative to sterile control. N=4/group. Combined with mRNA level analysis by qPCR of metabolic regulatory enzymes, in NTERT-1 keratinocytes, collected 6h post inoculation with *P. aeruginosa un*scratched or scratched. N=3/group. Data presented as log2 2^-ΔΔCt relative to control. To assess metabolic changes surrounding (A) L-arginine levels, and L-arginine metabolism; (B) *NOS2* mRNA and (C) L-citrulline levels; (D) *ARG1* mRNA and (E) L-ornithine and (F) urea levels; (G) agmatine levels. And downstream metabolism; (H) *AGMAT* and (I) *ODC* mRNA and (J) putrescine levels; (K) *AMD1* mRNA and (L) spermidine, (M) N8-acetylspermidine, (N) N1-8-diacetylspermidine, (O) spermine and (P) N1-acetylspermine levels; (Q) *PYCR3* mRNA and (R) L-proline levels. Individual points graphed with mean +/- SEM; p<0.05=*, p<0.01=**, p<0.001=***, p<0.0001=****. See also Figure S2.

The enzyme agmatinase (AGMAT) converts agmatine to putrescine and urea ^9^; L-ornithine metabolism also leads to putrescine production via ODC1. The addition of *P. aeruginosa* in keratinocyte cultures significantly increased keratinocyte mRNA levels of *AGMAT* (Figure 2h) and *ODC1* (Figure 2i), associated with higher putrescine (Figure 2j). Putrescine can be metabolised to the polyamines spermidine and spermine which is dependent on AMD1. In line with this, *AMD1* mRNA levels were increased by *P. aeruginosa* and scratch (Figure 2k); however, there was little change in polyamine levels (Figure 2l-p). L-ornithine metabolism can alternatively lead to L-proline production, via ornithine aminotransferase and pyrroline-5-carboxylate reductase ^9^, encoded by the gene *PYCR. PYCR3* was upregulated, in response to *P. aeruginosa* and in combination with scratch (Figure 2q); however, L-proline levels were unaltered (Figure 2r). We also observe reduced levels of L-asparagine (Figure S2c), L-glutamine (Figure S2d) and L-cysteine (Figure S2e) in wounded keratinocytes under *P. aeruginosa* challenge.

### *P. aeruginosa* enhances putrescine and polyamine acetylation in ARG1 deficient wounded keratinocytes

The heightened host cell gene regulation of L-arginine metabolism, induced by *P. aeruginosa,* was not seen in scratched keratinocytes in which ARG was knocked down by shRNAs (Figure 3). In these *shARG1* knockdown cells, mRNA levels of all metabolic enzymes analysed were reduced (Figure 3a-c, e-f), except for *ODC*, which remained relatively unchanged (Figure 3d). A similar reduction was also observed with nor-NOHA inhibition of arginase (Figure S3 a-f). The levels of the metabolites tested remained relatively unchanged in ARG1-deficient cells compared to infected control scratches (Figure S3 g-n), except for higher levels of putrescine (Figure 3g). Although polyamine levels remained constant (Figure 3h-i), there was a trend of higher acetylated spermidine (Figure 3j) and significantly higher levels of acetylated spermine (Figure 3k) in wounded infected keratinocytes on ARG1 knockdown (Figure S3g), suggesting increased polyamine flux. Alternative L-arginine metabolism is suggested by a significant reduction in L-glycine (Figure 4l). Of note, *P. aeruginosa* possesses genes to metabolise agmatine to putrescine (aguA, aguB); ornithine to putrescine (speC, speF); and putrescine to spermidine (speE) (Table S1, 2). This may suggest that, with a decline in host metabolism of the L-arginine pathway genes, *P. aeruginosa* could contribute to the maintenance or promotion of metabolite levels within keratinocytes.

**Figure 3.**
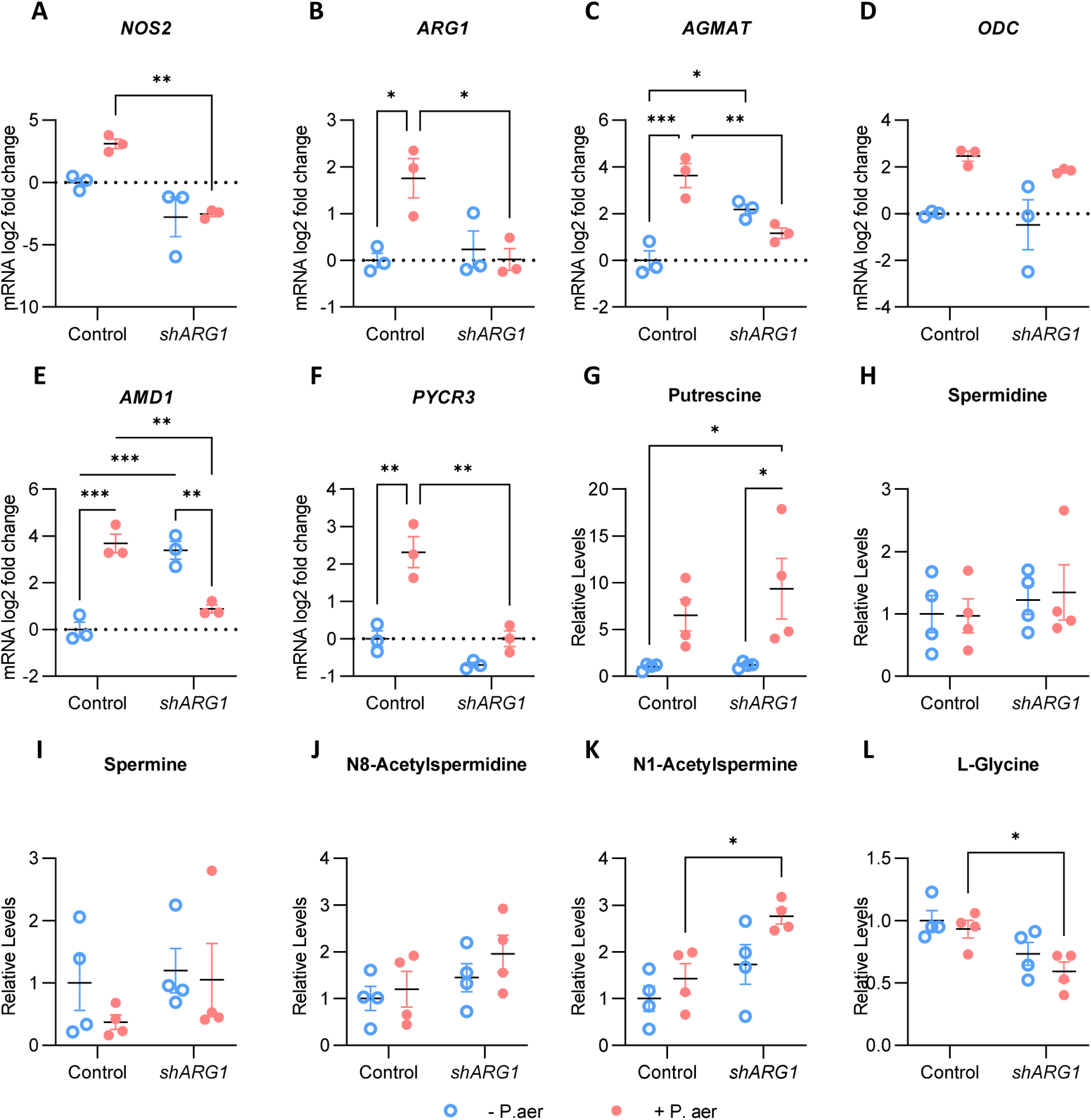
Wounded ARG1 deficient keratinocytes fail to promote host L-arginine metabolic activity in response to *P. aeruginosa;* however, exhibit increased polyamine metabolism. mRNA level analysis by qPCR of (A) *NOS2*, (B) *ARG1*, (C) *AGMAT*, (D) *ODC1*, (E) *AMD1* and (F) *PYCR3,* in *shARG1* and control cells 6h post scratch and co-culture with *P. aeruginosa*. N = 3/group. Data presented as log2 2^-ΔΔCt relative to control. Quantification of polyamine levels (G) putrescine, (H) spermidine, (I) N8-acetylspermidine, (J) spermine and (K) N1-acetylspermine and (L) amino acid L-glycine levels in scratched *shARG1* v control keratinocytes, 12h post co-culture with *P. aeruginosa*, determined by LC-MS. Data presented as metabolite levels relative to sterile control scratch. N=4/group. Individual points graphed with mean +/- SEM; p<0.05=*, p<0.01=**, p<0.001=***, p<0.0001=****. See also Figure S3.

**Figure 4.**
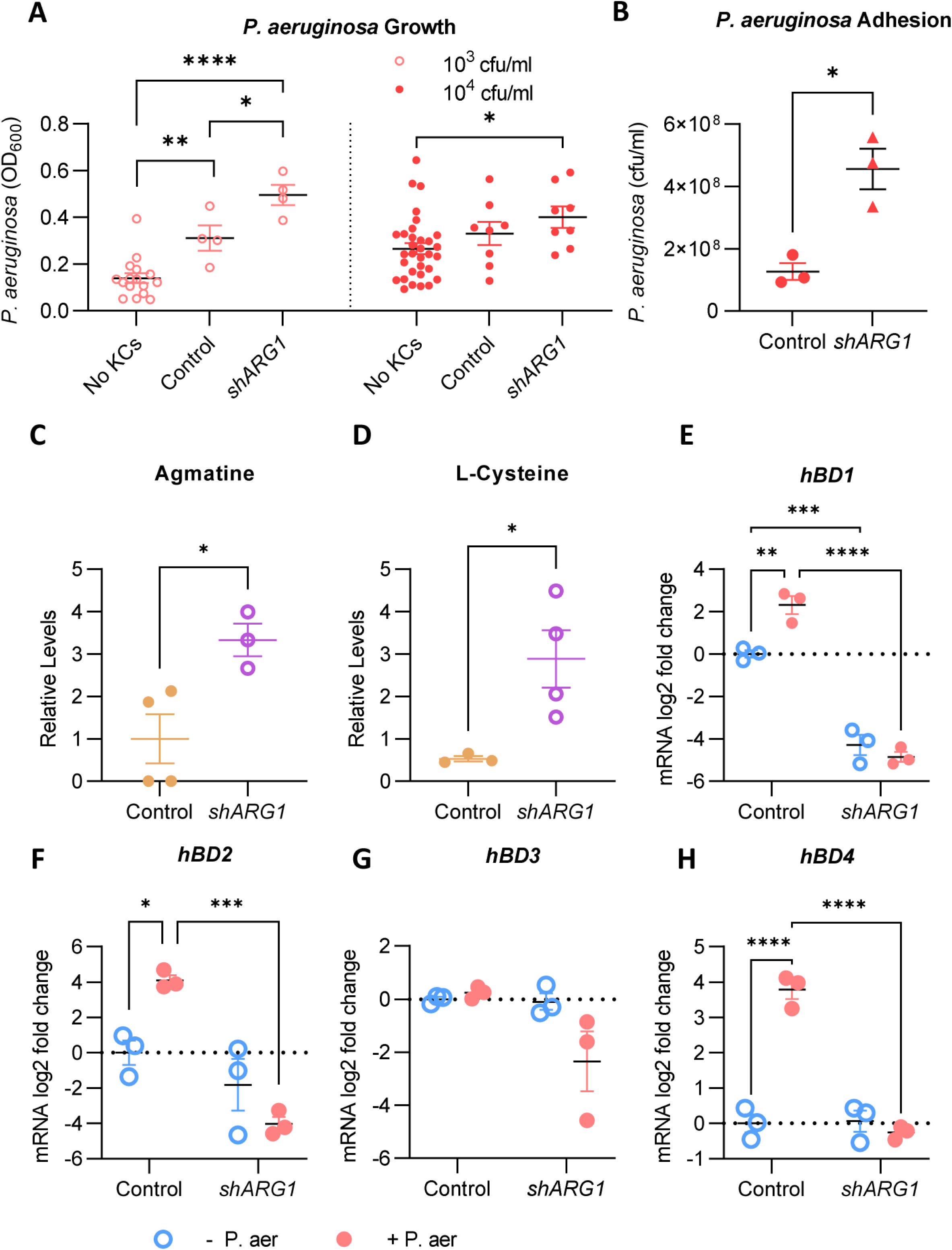
Host ARG1 deficiency in keratinocytes leads to enhanced bacterial growth by reduced AMP production. (A) Quantification of bacteria density determined by OD_600_ measurements, of *P. aeruginosa* 24h post co-culture with *shARG1* vs control NTERT-1 keratinocytes; n= 4-32/group. (B) *P. aeruginosa* adhesion to *shARG1* vs control NTERT-1 keratinocytes, determined by plate colony counts, expressed as CFU/ml; n=3/group. Quantification of (C) agmatine and (D) L-cysteine levels in *P. aeruginosa*, 12h post co-culture with scratched control or *shARG1* keratinocytes, determined by LC-MS. Data presented as metabolite levels relative to control. N=4/group. Analysis of *hBD1* (E), *hBD2* (F), *hBD3* (G), *hBD4* (H) mRNA levels in *shARG1* and control cells 6h post scratch and co-culture with *P. aeruginosa,* determined by qPCR, presented as log2 2^-ΔΔCt relative to control; n=3/group. All data presented as individual points graphed with mean +/- SEM; p<0.05=*, p<0.01=**, p<0.001=***, p<0.0001=****. See also Figure S4.

*P. aeruginosa* challenge strongly induced keratinocyte L-arginine metabolism upon wounding. However, in ARG1-depleted wounds, a higher level of polyamine metabolism was observed. When polyamine levels are high they can be acetylated, which can lead to export out of the cell ^32^; therefore, such increased levels of acetylated polyamines may support higher levels of secreted polyamines that could in turn influence *P. aeruginosa*.

### Host ARG1 is required to control bacterial growth via polyamines and AMPs

Bacterial load was analysed in bacteria-keratinocyte co-cultures. *P. aeruginosa* growth was increased when cultured with control keratinocytes and further enhanced in the presence of *shARG1* keratinocytes (Figure 4a, S4a), or when treated with nor-NOHA (Figure S4b). Consistent with this, colony counts of *P. aeruginosa* adhering to keratinocytes after 24h co-culture, were significantly higher in *shARG1* compared to controls (Figure 4b, S4c). The depletion of host ARG1 had little impact on polyamine levels within extracellular *P. aeruginosa* (Figure S4d-f); however, increased agmatine was observed (Figure 4e). Polyamine acetylation is also associated with polyamine catabolism by polyamine oxidases, which leads to release of H_2_O_2_. L-cysteine metabolism has been shown to exhibit anti-oxidative properties in bacteria. Higher L-cysteine is observed in *P. aeruginosa* (Figure 4f).

Keratinocytes detect and respond to the presence of microbes via production of anti-microbial peptides (AMPs), and the beta-defensins1-4 (hBD1-4) are known to be effective against *P. aeruginosa* ^33–35^*. hBD1, 2 and 4* mRNA levels were upregulated in keratinocyte scratches when challenged with *P. aeruginosa.* In contrast, *shARG1* keratinocytes (Figure 4g-j) had reduced *hBD1-4,* similarly reduced in nor-NOHA treated cells (Figure S4g-i).

## DISCUSSION

ARG expression by keratinocytes is required for effective cutaneous healing ^8,32^ and lower levels of epidermal ARG1 have been reported in non-healing DFUs ^8^. To determine whether reduced ARG1 in human aged skin and non-healing wounds impacts the microbial influence on repair, we used *in vivo* murine and *in vitro* keratinocyte co-culture models with the wound pathogen *P. aeruginosa.* We found inhibition of ARG impaired healing to a greater degree in response to bacterial challenge, associated with reduced keratinocyte proliferation, increased cell damage and inflammation. These functional impacts could be attributed to altered host metabolism in ARG1 deficient scratches, such as reduced *NOS2* (NO), L-glycine and heightened polyamine acetylation/export. These data also indicate shared metabolism and *P. aeruginosa* manipulation of the L-arginine pathway to evade host defences, supported by increased *P. aeruginosa-*derived agmatine and L-cysteine, and the increase in host putrescine and polyamine acetylation. Notably keratinocyte defence via AMPs was also reduced in *P. aeruginosa*-challenged scratches, collectively allowing increased *P. aeruginosa* growth, demonstrating the importance of ARG1 in keratinocyte microbial defence during wound repair.

We and others found that *P. aeruginosa* expresses multiple enzymes involved in L-arginine metabolic pathways (Figure S1a, 2b and Table S1, 2) ^31,36^, which suggests potential cross-utilisation of L-arginine pathway metabolites, leading to changes that can negatively influence healing outcome. *P. aeruginosa* colonisation of wounded keratinocytes was linked with enhanced host L-arginine metabolic activity, depleted stores of L-arginine, and increased levels of host regulatory enzymes (Figure 2), in line with infection of other cell types^37–40^, suggesting that infection contributes to upregulated L-arginine metabolism upon wounding ^8,13^. L-arginine metabolism by *P. aeruginosa* ^24,25^ (Figure S1a, Table S1, 2), could contribute to depletion of L-arginine and the increased L-citrulline, agmatine and putrescine observed in keratinocytes (Figure 2, 3, S3). Indeed, increased levels of the L-arginine derived amine, agmatine, in *P. aeruginosa* co-cultured with ARG1 deficient wounded keratinocytes supports that there is bacterial metabolism of L-arginine (Figure 4e). This also supports the idea of competition between host and bacteria over the L-arginine substrate, whereby in ARG1 depleted keratinocytes, L-arginine would be more ‘available’ for bacterial metabolism. Moreover, the higher putrescine and acetylated polyamine levels in infected *shARG1* keratinocyte scratches, supports the capacity of *P. aeruginosa* to promote polyamine metabolism.

Our results, showing the importance of ARG1 in controlling excessive inflammation and damage in response to bacterial infection, are in line with other studies investigating ARG in other disease models and cell types such as macrophages ^37,38,40^; however, inferred metabolic mechanisms differ to our findings. ARG1 deficiency, in *P. aeruginosa-*challenged keratinocytes, dampened host regulation of L-arginine metabolism upon wounding (Figure 3). Although we might expect the lack of functional ARG1 to result in no upregulation of ARG-directed metabolism (*ODC* > *AMD1* and *PYCR3*; Figure 3d-f), it was notable that L-arginine associated enzymes, *NOS2* and *AGMAT* were also lower (Figure 3a, c). This failure to upregulate host L-arginine metabolism could contribute to worsened healing responses, for example, via reduced NO, required for keratinocyte proliferation ^41^ and protection against apoptosis ^42^. Similarly, reduced metabolism of agmatine, and the increase in *P. aeruginosa* derived agmatine, could also negatively impact host cell proliferation ^43^. However, despite reduced host regulation by *shARG1* keratinocytes, L-arginine associated metabolite levels were maintained or even enhanced in the case of putrescine and acetylated polyamines, especially spermine. One possibility for this observation is that the reduced L-glycine observed in *P. aeruginosa* challenged *shARG1* scratches is due to AGAT/*GATM* converting L-arginine and L-glycine to guanidinoacetate and L-ornithine ^9^, thus maintaining L-ornithine levels. Our observation of heightened putrescine and acetylated spermine, suggests heightened polyamine metabolism, driven by *P. aeruginosa*, which is also reflected in other studies demonstrating promotion of polyamines by bacteria/microbiota ^44^, and could be considered beneficial for both bacteria and host. Intracellular levels of polyamines, spermidine and spermine, are maintained within a relatively constant range (Figure 3) which, can be achieved via biosynthesis, transport and catabolism. Acetylation is the first step in polyamine catabolism, rendering polyamines susceptible to either extracellular export or oxidation to lower polyamines by acetylpolyamine oxidase. However, oxidation reactions produce harmful intermediates, such as H_2_O_2_, which can lead to cell toxicity ^45^, and this may contribute to our observation of increased keratinocyte death. Collectively, altered metabolites could contribute to the reduced re-epithelialisation observed, both *in vitro* and *in vivo,* and the damaging cell responses in infected ARG-deficient wounds. This is in keeping with the fact that ARG1 levels can fall in ageing and when inflammation is protracted, rendering wounds less able to combat infection and heal swiftly ^46^. Our data supports these observations suggesting that common wound bacteria, such as *P. aeruginosa,* are able to scavenge and metabolise host L-arginine, contributing further to poor healing outcomes.

In contrast to other studies ^38,40^, which could be a reflection of the cell type, disease model and / or microbiota investigated, we found a host keratinocyte ARG deficiency promoted *P. aeruginosa* growth (Figure 4a, b). Akin to this, non-healing DFUs, even if not clinically infected, have a higher bacterial burden ^47^, including typical wound pathogens, such as *P. aeruginosa* ^26^, shown to impair healing ^48,49^. In our *in vitro* model, the heightened bacterial burden was linked with higher keratinocyte cell death (Figure 1). Epithelial cell death can result in the release of additional factors that support bacterial growth ^50^; therefore, increased scavenging of L-arginine-derived host metabolites by bacteria may contribute to the higher bacterial burden we saw.

L-arginine metabolism has multiple impacts on bacteria via downstream metabolites including NO ^51–53^. We observed increased *NOS2* mRNA in *P. aeruginosa* infected keratinocytes which could promote NO production ^9^. In response, *P. aeruginosa* could preferentially metabolise L-arginine to produce agmatine (Figure 4), which can inhibit NOS activity ^10,54,55^. However, host cells may circumvent this by metabolising agmatine via AGMAT, shown in lung epithelial cells to enhance NOS2 and NO production ^56^. Indeed, we found, reduced *NOS2* and *AGMAT* in *shARG1* keratinocytes, suggesting inhibition of NOS activity and less NO production that may contribute to higher bacterial load. The suggested export of polyamines from host cells could also affect bacteria. Polyamines may promote *P. aeruginosa* growth and survival ^25,57,58^; however, intracellular levels of polyamines in *P. aeruginosa* were unaltered. As previously mentioned, polyamine levels are tightly regulated to prevent harmful toxic effects induced by high levels of polyamine conversion; the increased agmatine in *P. aeruginosa* co-cultured with *shARG1* keratinocytes (Figure 4e) could support reduced polyamine biosynthesis, due to increased availability and import via *P. aeruginosa* polyamine transporters ^59,60^, less demanding than biosynthesis. Furthermore, our observations of heightened agmatine and L-cysteine (Figure 4e-f), that are known to benefit bacterial growth and defence mechanisms ^54,55,61–63^, may also support *P. aeruginosa* growth. There were other consequences of ARG1 deficiency on wound infection. ARG1 was required for the induction of *β-defensins 1-4* (*hBD1-4*) and other data from our lab has supported the impact of ARG deficiency on AMP expression ^8,64^. Therefore, the lack of AMP induction in *shARG1* keratinocytes is another contributing factor in the failure to control *P. aeruginosa* growth. Collectively, these metabolic changes in host and *P. aeruginosa,* together with reduced anti-microbial products, suggest that bacteria can actively benefit from scavenging available host products of L-arginine metabolism and evade host keratinocyte defence mechanisms. Moreover, AMPs are pleiotropic molecules that can also affect cellular migration and proliferation ^65,66^. As such, reduced *hBDs* may have directly impacted the reduced re-epithelialisation observed in ARG deficient wounds (Figure 1).

This study demonstrates the importance of ARG1 in regulating the keratinocyte wound healing response and controlling microbial-induced inflammation and cell damage. We suggest that the combinatorial impact of microbial influence, compounded by ARG1 impairment, contributes to the poor healing outcome in chronic wounds. The L-arginine metabolic pathway encompasses numerous biological functions across human and microbial species, and modulation of this pathway requires further examination in the context of host microbiota interactions in cutaneous disease. Specific targeting of this metabolic network could prove beneficial in controlling both infection and inflammation related skin conditions and repair.

## Acknowledgments

The work was funded by a BBSRC PhD awarded to SC, AJM and RC. Metabolite analysis was funded by the A*STAR Skin Research Laboratories and performed by the Skin Metabolomics Platform. The Bioimaging Facility microscopes used in this study were purchased with grants from BBSRC, Wellcome and the University of Manchester Strategic Fund. Special thanks go to Peter March and Steve Marsden for their help with the microscopy.

## Author contributions

R.C. conducted methodology, formal analysis, and wrote the original draft; S.C. and A.J.M conceptualised, supervised the work, acquired findings and reviewed the draft; C.O. provided resources, funded the work, supervised and reviewed the draft; D.S. and C.D. conducted methodology; H.T. provided resources and methodology; S.H.L and Y.P.L. conducted methodology and analysis; L.V. provided resources, conducted formal analysis and reviewed the paper. All authors contributed to writing the manuscript text.

## Materials availability

This study did not generate new unique reagents.

## Data availability

Data in this study can be shared by the lead contact upon request after publication

## METHODS

### Murine Wounding

Animal studies were conducted according to the Animals (Scientific Procedures) Act 1986 under project license P8721BD27 approved by the UK Home Office. 6-week-old female C57BL/6J wild type mice were purchased from Envigo (Oxfordshire, UK) delivered to the University of Manchester’s Biological Services Facility and acclimatised for 2 weeks prior to experimental use. Mice were housed in isolator cages with ad libitum food and water. The room was maintained at a constant temperature of 21°C, with 45-65% humidity on a 12-hour light-dark cycle. Mice were subcutaneously injected with 50µl nor-NOHA (400µg/ml) or PBS as vehicle alone at the site of wounding 24 and 2hrs prior to wounding. Mice were anaesthetised by isoflurane inhalation (induced at 2L/min and maintained at 0.5L/min) and buprenorphine (0.1mg/kg) administered subcutaneously as analgesia. The posterior aspect of each mouse was shaved and sanitised using ethanol. Using a biopsy punch, two equidistant 6mm full-thickness dorsal excisions were made through both skin and panniculus carnosus muscle ^67,68^. Wounds were inoculated with 20µl containing 10^2^ cfu *P. aeruginosa* or saline (0.9% NaCl) as vehicle alone, topically applied to the wound bed. Sterile 1cm^2^ dressings, pre-moistened with saline to maintain a moist wound environment, were applied to each wound, covered with 3M^TM^ Tegaderm^TM^ Film (3M, Bracknell, UK), and secured with Mastisol^®^ Liquid Adhesive (Eloquest Healthcare, Ferndale, MI) applied to the edges. Following wounding, all mice were individually housed with sizzle nest bedding only (no sawdust); provided with mash as well as their normal dry pellets and water and temporarily housed in warm ScanTainers until alert and fit to return. Wounds were left to heal by secondary intention. Further treatments of nor-NOHA or vehicle were injected through the wound adhesive applied to the wound site daily until sacrificed 3 days post wounding by a rising concentration of CO_2_ followed by dislocation of the neck and tissue collected. Wounds were excised down to the underlying muscle and normal skin taken from the dorsal shoulder region. Wounds were bisected and fixed in formalin, before embedding in paraffin wax for histological analysis; snap frozen in liquid nitrogen and stored at −80°C for biochemical analysis.

### Histology - Haemotoxylin and Eosin (H&E) Staining

Formalin fixed (1.075M 37% Formaldehyde, 0.35M Acetic Acid, 0.155M NaCl, 1.38nM hexadecyl trimethyl-ammonium bromide in dH2O) samples were processed (Microm Spin STP120 tissue processor (ThermoFisher Scientific, Loughborough, UK)) and wax embedded (Embedding Station; Leica Biosystems, Newcastle Upon Tyne, UK) ensuring the bisected edge (the wound midline) was faced down to provide a full skin wound profile to include the epidermis and dermis. 5µm thick tissue sections were obtained using a manual rotary microtome (Leica Biosystems) and adhered to Vectabond (Vector Laboratories, Peterborough, UK) coated glass slides. Sections were H&E stained using Shandon Instant Hematoxylin (ThermoFisher Scientific) and 1% Eosin (Sigma, Gillingham, UK). Coverslips were mounted using Pertex mounting media (CellPath, Newtown, UK). Images were acquired using a Nikon Eclipse Ci microscope using x4/0.13 Plan Fluor objective, with a Nikon DS-Fi3 camera and NIS-Elements software (Nikon, Kingston Upon Thames, UK). Quantification of wound measurements were performed manually using Image Pro Premier software v9.3 (Media Cybernetics, Abingdon, United Kingdom), analysed for wound area, wound width and percent re-epithelialisation. The wound area was considered the area of granulation tissue beneath the clot to the margins of normal skin on either side of the wound. Percentage re-epithelialisation was determined by dividing the sum length of each neo-epidermal tongue by the total distance required to fully close the wound.

### Bacterial Culture

*P. aeruginosa* (NCTC-10781, alternative resource assignments ATCC 23267; CCTM A18; GESSARD M5; KM 256; RH 2583), sourced from a human leg ulcer, was purchased from the National Collection of Type Cultures (NCTC) of Public Health England (Salisbury, UK). *Staphylococcus epidermidis* (DSM20042), *S. capitis* (DSM20325) and *S. hominis* (DSM20328), purchased from the Leibniz Institute DSMZ - German Collection of Microorganisms and Cell Cultures GmbH (Braunschweig, Germany). Stocks were revived from cryobeads (Microbank ™ cryovials; Pro-lab Diagnostics, Cheshire, UK) streaked across solid tryptone soy yeast extract agar plates (TSYEA: 3% tryptone soy 0.3% yeast, 1.5% agar, pH7.2 (ThermoFisher Scientific, Loughborough, UK)), grown in aerobic conditions at 37°C for 24-48hrs. 10ml TSYE broth (TSYEB: 3% tryptone soy, 0.3% yeast, pH7.2 (ThermoFisher Scientific)), inoculated with approx. five single colonies were cultured overnight in aerobic conditions at 37°C with shaking for experimental use. Bacterial concentration was determined by OD_600_ quantification of liquid cultures and corresponding colony growth of bacterial dilutions. For experimental use overnight cultures were adjusted to 10^7^ cfu/ml and further diluted to the required concentration.

### Keratinocyte Culture

The human immortalised keratinocyte HaCaT cell line (T0020001; Addex Bio, Caltag Medsystems Ltd, UK) was routinely cultured in DMEM (Gibco, UK) supplemented with 10% Foetal Bovine Serum (Gibco) and Sodium Pyruvate (final concentration of 2mM; Sigma-Aldrich, Gillingham, UK), at 37°C with 5% CO_2_. No antibiotics were used during routine culture or experimental use.

N/TERT-1 ^69^ cells were cultured at 37°C with 5% CO_2_ in Keratinocyte Serum-Free Media (K-SFM) (Gibco) supplemented with 0.2ng/ml Epidermal Growth Factor (EGF) (Gibco), 25μg/ml Bovine Pituitary Extract (BPE) (Gibco) and 0.3mM CaCl_2_ (PromoCell GmbH, Heidelberg, Germany), as described previously (Rheinwald *et al.*, 2002). NTERT-1 cells were transferred to a defined keratinocyte media, DFK-2, for experimental use, which consists of a 1:1 ratio of DFK-1 media [1:1 high glucose, calcium and glutamine free DMEM (Gibco) : Ham’s F-12 (Gibco) OR F-12 (1x +GlutaMax Tm/ Nutrient Mixture (Ham) (Gibco), supplemented with 0.2 ng/ml EGF, 25 μg/ml BPE and 1.5 mM L-glutamine (Gibco) and complete K-SFM ^70^.

Generation of shARG1 knockdown keratinocytes, by shRNA transfection of NTERT-1 cells, was described previously with validation of knockdown efficacy shown ^64^. shARG1 and scrambled control were maintained in NTERT-1 media with puromycin (1μg/ml, Gibco) selection. Puromycin was excluded during experimental use.

### Keratinocyte Scratch Assays

Keratinocyte scratch assays were employed to assess the impact of bacterial presence on keratinocyte proliferation and migration. HaCaTs or N/TERT-1 cells were seeded at a density of 10^4^ cells/well of an Incucyte® Imagelock 96-well Plate (Sartorius, Germany). Plates were incubated under standard culture conditions for 3 days until confluent. Monolayers were stained with 2µM CellTrackerTM Red CMTPX Dye (Invitrogen, UK) for 50min, washed with sterile PBS, scratched with an Incucyte® Woundmaker Tool (Sartorius) and rinsed with sterile PBS to remove cell debris. Overnight bacterial cultures were diluted to 10^7^ cfu/ml, then serially diluted 10-fold to 10^3^ cfu/ml in standard HaCaT or DFK-2 NTERT-1 media. 100µl inoculated or sterile media was added and incubated at 37°C for 24hrs. Images were acquired every 2hrs on an Incucyte S3 Live Cell Analysis system using a *10x/ 1.22 Plan Fluor* objective and the red filter set for fluorescent images (Sartorius). Images were measured manually using Image Pro Premier software (Media Cybernetics, Abingdon, United Kingdom) and the degree of wound closure as a percentage was calculated based on scratch area after specified duration (D), in relation to area at time zero (T0) using the equation ((T0-D)/T0)100.

### Keratinocyte Viability and Proliferation by Live / Dead Counts

Keratinocytes were seeded at 10^5^ cells/well into 6-well culture plates and grown to approx. 60% confluency. Overnight bacterial cultures were diluted to 10^7^ cfu/ml and serially diluted 10-fold to 10^3^ cfu/ml in keratinocyte media. 2ml of inoculated or sterile media was added and co-cultures were incubated for 24hrs at 37°C 5% CO_2_. Culture medium was set aside to retain any dead cells from each well and keratinocytes were washed in sterile PBS, trypsinised to detach and neutralised in media containing FBS. Cells were pelleted by centrifugation (300g 5min) and supernatant removed. Pellets were resuspended in the original culture medium, and cell counts, and viability determined by AO/PI fluorescent staining to identify viable (green) / non-viable (red) keratinocytes respectively, quantified using the Cellometer Auto 2000 automated cell counter (Nexcelom, Manchester, UK). Total, live and dead cell counts, and cell diameter was obtained, and percentage viability calculated.

### Apoptosis / Necrosis of Keratinocytes by Flow Cytometry

NTERT-1 shARG and control keratinocytes were seeded at 10^5^ cells/well into 6-well culture plates and grown to approx. 60% confluency and co-cultured with live P. aeruginosa for 24hrs at 37°C 5% CO_2_. Cells were trypsinised and washed twice with cold PBS then resuspended in 1ml 1x Annexin-V binding buffer (10 mM HEPES/NaOH, 140 mM NaCl, 2.2 mM CaCl2, pH 7.4; BD Bioscience, Cambridge, UK). Cells were stained with AF647-Annexin-V (BioLegend, UK) for 30 min on ice, diluted to approx. 10^5^ cells/500ul. 0.25 µg/ml of 4′, 6-diamidino-2-phenylindole (DAPI) (Cell Signaling Technology, Leiden, NL) was added just prior to quantification using the BD LSRII (BD Bioscience). Total cells were gated using forward scatter (FSC) versus side scatter (SSC), from which a single cell population was gated using FSC-area (A) versus FSC-height (H). Single stain controls for apoptosis pre-treated with 15mM camptothecin for 5-6hrs stained with Annexin-V; necrosis pre-heated for 10sec in the microwave stained with DAPI; and unstained sterile keratinocytes were used to aid acquisition parameters and gating strategies, applied to all technical repeats to ensure consistency. DAPI against Annexin-V scatter plots were created from the single cell population and quadrants applied to provide % viable, early apoptotic, late apoptotic and necrotic cells, analysed using FlowJo software (Tree Star© V10.5.3)

### Reverse Transcriptase Real Time PCR

Keratinocytes were seeded at 10^5^ cells / well of 6 well tissue culture plates and grown to full confluency. Cells were scratched and inoculated with live bacteria and incubated at 37C 5% CO_2_ for 6 or 24hrs post treatment and collected for RNA extraction. Total RNA was isolated from keratinocytes by TRIzol (Invitrogen, Paisley, UK) / chloroform extraction and column-based purification using the Purelink RNA mini kit (Invitrogen) according to the manufacturer’s instructions and quantified using a nanodrop (Invitrogen). cDNA was transcribed from 1μg of RNA using the GoScript RT kit (Promega, Madison, WI) and quantitative PCR performed using the PowerUp Sybr Green Master Mix (Thermo Fisher Scientific) and LightCycler 480 Instrument (Roche, Welwyn Garden City, UK). Primers were designed using PrimerBlast. For each primer set, melt curves were used to determine amplification specificity and serial dilutions used to determine primer efficiency. Each sample was performed in duplicate, and relative expression was determined using the delta-delta ct method. CT values normalised to the housekeeper YWHAZ. The full primer sequences are listed in Table S3.

### LC-MS Metabolite Analysis of Keratinocytes and *P. aeruginosa* in Co-cultures

#### Sample processing of keratinocytes and *P. aeruginosa*

2×10^6^ keratinocytes were seeded into 10 cm tissue culture dishes and grown in DFK2 until confluent. Monolayers were scratched vertically and horizontally over 30× using a sterile 1 ml tip. Overnight *P. aeruginosa* cultures were diluted to 10^3^ cfu/mL in DFK2 and 10 mL of inoculated or sterile media was added and incubated at 37°C 5% CO_2_. After 12h, unattached bacteria were collected and filtered using a 5 µm pore size filter. Keratinocytes were washed 3× and detached with trypsin. Bacteria and keratinocytes were pelleted, washed twice and cell pellets, containing 10^7^ or 10^6^ cells respectively, were stored at −80°C prior to analysis.

For polyamines analysis, a volume of 100 µL of acetonitrile:water (50:50) containing internal standards (100 ng/mL of D^8^-putrescine (DLM-6573, Cambridge Isotope Laboratories, Andover, MA)) was added to each sample. The samples were subjected to three cycles of freeze-thaw and ultrasonication cell lysis. Each cycle involved placing the samples on dry ice for 30 minutes, followed by thawing at room temperature, vortexing, and ultrasonication for 1 minute. Samples were then centrifuged at 18,000 *g* for 10 min at 4°C, and 50 µL of the supernatants were obtained. Samples were sent for LC-MS analysis at 0.2 µL injection volume. For amino acid analysis, a volume of 20 µL of acetonitrile:water (50:50) containing internal standards (30 ng/mL of D^4^-lysine (DLM-2640, Cambridge Isotope Laboratories)) was added to each sample. The samples were subjected to three cycles of freeze-thaw and ultrasonication cell lysis. Each cycle involved placing the samples on dry ice for 30 minutes, followed by thawing at room temperature, vortexing, and ultrasonication for 1 minute. Samples were then centrifuged at 18,000 *g* for 10 min at 4°C, and 10 µL of the supernatants were obtained. Keratinocytes samples were diluted by 20x prior to analysis at 1 µL injection volume, while *P. aeruginosa* samples were sent for analysis at 2 µL injection volume.

#### Liquid chromatography mass spectrometry

Multiple reaction monitoring was performed on a Thermo Scientific Vanquish Duo UHPLC coupled to a Quantis TSQ Triple Quadrupole Mass Spectrometer. Mass spectrometry was operated in electrospray ionization (ESI) positive mode, with source parameters as follows: spray voltage 3.5 kV (positive), sheath gas 50 [arbitrary units], auxiliary gas 18 [arbitrary units], sweep gas 0 [arbitrary units], ion transfer tube temperature 325°C, vaporizer temperature 230°C.

For polyamines analysis, chromatographic separation was done on a Waters Atlantis Premier BEH Z-HILIC column (1.7 µm, 2.1 mm × 100 mm) maintained at 45°C. Mobile phase A comprises 10 mM ammonium formate at pH 3, while mobile phase B comprises 0.1% formic acid in acetonitrile. Chromatographic separation was achieved at 0.35 mL/min using the gradient elution as follows: 0.0 min 65%B, 0.5 min 65%B, 8.0 min 30%B, 12.0 min 30%B, 12.1 min 65%B, 15.0 min 65%B. For amino acids analysis, chromatographic separation was done on a Waters Acquity HSS T3 column (1.8 µm, 2.1 mm × 100 mm) maintained at 45°C. Mobile phase A comprises 0.1% formic acid in water, while mobile phase B comprises 0.1% formic acid in acetonitrile. Chromatographic separation was achieved at 0.3 mL/min using the gradient elution as follows: 0.0 min 0%B, 2.0 min 100%B (curve 9), 6.0 min 100%B, 6.1 min 0%B, 10.0 min 0%B. Details on compound-dependent parameters are described in the Table S4. Peak area integration and data analysis were performed on Skyline, and metabolites’ areas were normalised to the internal standards and presented relative to the control group.

### Bacterial Growth and Adhesion in Keratinocyte Co-cultures

Keratinocytes were seeded into 96-well tissue culture plates and grown under standard culture conditions until confluent. Overnight bacterial cultures were diluted to 10^7^ cfu/ml and serially diluted to 10^4^, 10^3^ cfu/ml in phenol red free DFK2. Keratinocytes were washed with sterile PBS, 200µl of inoculated or sterile media was added and incubated at 37°C 5% CO2. After 24hrs bacterial growth was determined by OD_600_ measurements (VersaMax microplate reader; Molecular Devices LLC, San Jose, CA), blanked against wells containing keratinocytes and sterile media. Media was removed and wells washed to remove non-adherent bacteria. Keratinocytes were lysed in 100 µl 0.1% Triton-X-100 for 10 minutes at RT and bacterial CFU determined by serial dilution and plating onto agar.

### Bacteria Whole Genome Sequencing

Complete Genome Sequencing and Analysis was performed by BMKgene (Biomarker Technologies (BMK) GmbH, Munster, Germany). For genome assembly, the filtered reads were assembled by Spades v3.6.2 software. For genome component prediction, Coding genes prediction was performed by Prodigal v2.6.3. The GenBlastA v1.0.4 program was used to scan the whole genomes after masking predicted functional genes. Putative candidates were then analyzed by searching for non-mature mutations and frame-shift mutations using GeneWise v2.2.0. Transfer RNA (tRNA) genes were predicted with tRNAscan-SE v2.0, Ribosome RNA (rRNA) genes were predicted with Infernal v1.1.3. Repetitive sequences were predicted using RepeatMasker. PhiSpy v2.3 is used for prophage prediction and CRT v1.2 was used for CRISPR identification. IslandPath-DIMOB v0.2 was used to predict genomic island in genome. antiSMASH v5.0.0 was used to predict secondary metabolic gene clusters, and PromPredict v1 was used for promoter prediction.For functional annotation, the predicted protein was blast (e-value: 1e-5) against Nr,Swiss-Prot,TrEMBL,KEGG,eggNOG, Blast2go was used for GO annotation. Protein sequences of predicted genes were BLAST against sequences in KEGG database to obtain corresponding KO number and position in the pathway.

### Statistical Analysis

Gaussian distribution for all data was assessed statistically using Shapiro-Wilk normality test and visually via QQ plots. Ordinary or repeated measures two-way ANOVA followed by Sidak’s multiple comparisons test was performed for all grouped data with two factors and adjusted *P*-values were reported. If normality could not be assumed, statistical comparisons between groups were determined using the Mann‒Whitney test at each comparison set, and the two-tailed *P*-value was reported. Ordinary one-way ANOVA was used for single factor data with three or more groups followed by Tukey’s multiple comparisons, or Kruskal‒ Wallis test with Dunn’s multiple comparisons if normality could not be assumed. All statistical analysis was performed using GraphPad Prism 9, version 9.1.2 (GraphPad Software, San Diego, CA). A probability value of less than 0.05 was considered statistically significant.

## References

1. Falanga, V., Isseroff, R.R., Soulika, A.M., Romanelli, M., Margolis, D., Kapp, S., Granick, M., and Harding, K. (2022). Chronic wounds. Nat Rev Dis Primers 8, 50. 10.1038/s41572-022-00377-3.

2. Campbell, L., Saville, C.R., Murray, P.J., Cruickshank, S.M., and Hardman, M.J. (2013). Local arginase 1 activity is required for cutaneous wound healing. J Invest Dermatol 133, 2461–2470. 10.1038/jid.2013.164.

3. Dixit, R., Debnath, A., Mishra, S., Mishra, R., Bhartiya, S.K., Pratap, A., and Shukla, V.K. (2021). A Study of Arginase Expression in Chronic Non-healing Wounds. Int. J. Lower Extremity Wounds. 10.1177/15347346211012381.

4. Abd El-Aleem, S.A., Abd-Elghany, M.I., Ali Saber, E., Jude, E.B., and Djouhri, L. (2020). A possible role for inducible arginase isoform (AI) in the pathogenesis of chronic venous leg ulcer. J. Cell. Physiol. 235, 9974–9991. 10.1002/jcp.29812.

5. Abd-El-Aleem, S.A., Ferguson, M.W., Appleton, I., Kairsingh, S., Jude, E.B., Jones, K., McCollum, C.N., and Ireland, G.W. (2000). Expression of nitric oxide synthase isoforms and arginase in normal human skin and chronic venous leg ulcers. J Pathol 191, 434–442. 10.1002/1096-9896(2000)9999:9999<::AID-PATH654>3.0.CO;2-S.

6. Kampfer, H., Pfeilschifter, J., and Frank, S. (2003). Expression and activity of arginase isoenzymes during normal and diabetes-impaired skin repair. J Invest Dermatol 121, 1544–1551. 10.1046/j.1523-1747.2003.12610.x.

7. Miao, M., Niu, Y., Xie, T., Yuan, B., Qing, C., and Lu, S. (2012). Diabetes-impaired wound healing and altered macrophage activation: a possible pathophysiologic correlation. Wound Repair Regen 20, 203–213. 10.1111/j.1524-475X.2012.00772.x.

8. Crompton, R.A., Williams, H., Campbell, L., Hui Kheng, L., Saville, C., Ansell, D.M., Reid, A., Wong, J., Vardy, L.A., Hardman, M.J., and Cruickshank, S.M. (2022). An Epidermal-Specific Role for Arginase1 during Cutaneous Wound Repair. J Invest Dermatol 142, 1206–1216 e1208. 10.1016/j.jid.2021.09.009.

9. Morris, S.M., Jr. (2016). Arginine Metabolism Revisited. J Nutr 146, 2579S–2586S. 10.3945/jn.115.226621.

10. Morris, S.M., Jr. (2004). Enzymes of arginine metabolism. J Nutr 134, 2743S–2747S; discussion 2765S-2767S. 10.1093/jn/134.10.2743S.

11. Bailey, J.D., Diotallevi, M., Nicol, T., McNeill, E., Shaw, A., Chuaiphichai, S., Hale, A., Starr, A., Nandi, M., Stylianou, E., et al. (2019). Nitric Oxide Modulates Metabolic Remodeling in Inflammatory Macrophages through TCA Cycle Regulation and Itaconate Accumulation. Cell Rep 28, 218–230 e217. 10.1016/j.celrep.2019.06.018.

12. Yamamoto, N., Oyaizu, T., Enomoto, M., Horie, M., Yuasa, M., Okawa, A., and Yagishita, K. (2020). VEGF and bFGF induction by nitric oxide is associated with hyperbaric oxygen-induced angiogenesis and muscle regeneration. Sci Rep 10, 2744. 10.1038/s41598-020-59615-x.

13. Szondi, D.C., Wong, J.K., Vardy, L.A., and Cruickshank, S.M. (2021). Arginase Signalling as a Key Player in Chronic Wound Pathophysiology and Healing. Frontiers in Molecular Biosciences 8, 773866. 10.3389/fmolb.2021.773866.

14. Martí i Líndez, A.A., and Reith, W. (2021). Arginine-dependent immune responses. Cell. Mol. Life Sci. 78, 5303–5324. 10.1007/s00018-021-03828-4.

15. Piipponen, M., Li, D., and Landen, N.X. (2020). The Immune Functions of Keratinocytes in Skin Wound Healing. Int J Mol Sci 21. 10.3390/ijms21228790.

16. Kollisch, G., Kalali, B.N., Voelcker, V., Wallich, R., Behrendt, H., Ring, J., Bauer, S., Jakob, T., Mempel, M., and Ollert, M. (2005). Various members of the Toll-like receptor family contribute to the innate immune response of human epidermal keratinocytes. Immunology 114, 531–541. 10.1111/j.1365-2567.2005.02122.x.

17. Wanke, I., Steffen, H., Christ, C., Krismer, B., Gotz, F., Peschel, A., Schaller, M., and Schittek, B. (2011). Skin commensals amplify the innate immune response to pathogens by activation of distinct signaling pathways. J Invest Dermatol 131, 382–390. 10.1038/jid.2010.328.

18. Verbanic, S., Shen, Y., Lee, J., Deacon, J.M., and Chen, I.A. (2020). Microbial predictors of healing and short-term effect of debridement on the microbiome of chronic wounds. NPJ Biofilms Microbiomes 6, 21. 10.1038/s41522-020-0130-5.

19. Armstrong, D.G., Boulton, A.J.M., and Bus, S.A. (2017). Diabetic Foot Ulcers and Their Recurrence. N Engl J Med 376, 2367–2375. 10.1056/NEJMra1615439.

20. Qualls, J.E., Subramanian, C., Rafi, W., Smith, A.M., Balouzian, L., DeFreitas, A.A., Shirey, K.A., Reutterer, B., Kernbauer, E., Stockinger, S., et al. (2012). Sustained generation of nitric oxide and control of mycobacterial infection requires argininosuccinate synthase 1. Cell Host Microbe 12, 313–323. 10.1016/j.chom.2012.07.012.

21. Thurlow, L.R., Joshi, G.S., Clark, J.R., Spontak, J.S., Neely, C.J., Maile, R., and Richardson, A.R. (2013). Functional modularity of the arginine catabolic mobile element contributes to the success of USA300 methicillin-resistant Staphylococcus aureus. Cell Host Microbe 13, 100–107. 10.1016/j.chom.2012.11.012.

22. Kwon, D.H., and Lu, C.D. (2007). Polyamine effects on antibiotic susceptibility in bacteria. Antimicrob Agents Chemother 51, 2070–2077. 10.1128/AAC.01472-06.

23. Zhang, C., Hu, Z., Lone, A.G., Artami, M., Edwards, M., Zouboulis, C.C., Stein, M., and Harris-Tryon, T.A. (2022). Small proline-rich proteins (SPRRs) are epidermally produced antimicrobial proteins that defend the cutaneous barrier by direct bacterial membrane disruption. Elife 11. 10.7554/eLife.76729.

24. Rinaldo, S., Giardina, G., Mantoni, F., Paone, A., and Cutruzzola, F. (2018). Beyond nitrogen metabolism: nitric oxide, cyclic-di-GMP and bacterial biofilms. FEMS Microbiol Lett 365, fny029. 10.1093/femsle/fny029.

25. Hernandez, V.M., Arteaga, A., and Dunn, M.F. (2021). Diversity, properties and functions of bacterial arginases. FEMS Microbiol Rev 45, fuab034. 10.1093/femsre/fuab034.

26. Kalan, L.R., Meisel, J.S., Loesche, M.A., Horwinski, J., Soaita, I., Chen, X., Uberoi, A., Gardner, S.E., and Grice, E.A. (2019). Strain- and Species-Level Variation in the Microbiome of Diabetic Wounds Is Associated with Clinical Outcomes and Therapeutic Efficacy. Cell Host Microbe 25, 641–655 e645. 10.1016/j.chom.2019.03.006.

27. Pastar, I., Stojadinovic, O., Yin, N.C., Ramirez, H., Nusbaum, A.G., Sawaya, A., Patel, S.B., Khalid, L., Isseroff, R.R., and Tomic-Canic, M. (2014). Epithelialization in Wound Healing: A Comprehensive Review. Adv Wound Care (New Rochelle) 3, 445–464. 10.1089/wound.2013.0473.

28. Coleman, C.S., Hu, G., and Pegg, A.E. (2004). Putrescine biosynthesis in mammalian tissues. Biochem J 379, 849–855. 10.1042/BJ20040035.

29. Paulson, N.B., Gilbertsen, A.J., Dalluge, J.J., Welchlin, C.W., Hughes, J., Han, W., Blackwell, T.S., Laguna, T.A., and Williams, B.J. (2014). The arginine decarboxylase pathways of host and pathogen interact to impact inflammatory pathways in the lung. PLoS One 9, e111441. 10.1371/journal.pone.0111441.

30. Zhu, M.Y., Iyo, A., Piletz, J.E., and Regunathan, S. (2004). Expression of human arginine decarboxylase, the biosynthetic enzyme for agmatine. Biochim Biophys Acta 1670, 156–164. 10.1016/j.bbagen.2003.11.006.

31. Mercenier, A., Simon, J.P., Haas, D., and Stalon, V. (1980). Catabolism of L-arginine by Pseudomonas aeruginosa. J Gen Microbiol 116, 381–389. 10.1099/00221287-116-2-381.

32. Nakanishi, S., and Cleveland, J.L. (2021). Polyamine Homeostasis in Development and Disease. Med Sci (Basel) 9. 10.3390/medsci9020028.

33. Singh, P.K., Jia, H.P., Wiles, K., Hesselberth, J., Liu, L., Conway, B.A., Greenberg, E.P., Valore, E.V., Welsh, M.J., Ganz, T., et al. (1998). Production of beta-defensins by human airway epithelia. Proc Natl Acad Sci U S A 95, 14961–14966. 10.1073/pnas.95.25.14961.

34. Harder, J., Bartels, J., Christophers, E., and Schroder, J.M. (2001). Isolation and characterization of human beta -defensin-3, a novel human inducible peptide antibiotic. J Biol Chem 276, 5707–5713. 10.1074/jbc.M008557200.

35. Smiley, A.K., Gardner, J., Klingenberg, J.M., Neely, A.N., and Supp, D.M. (2007). Expression of human beta defensin 4 in genetically modified keratinocytes enhances antimicrobial activity. J Burn Care Res 28, 127–132. 10.1097/BCR.0b013E31802C88FD.

36. Lu, C.D. (2006). Pathways and regulation of bacterial arginine metabolism and perspectives for obtaining arginine overproducing strains. Appl Microbiol Biotechnol 70, 261–272. 10.1007/s00253-005-0308-z.

37. Gobert, A.P., Cheng, Y., Akhtar, M., Mersey, B.D., Blumberg, D.R., Cross, R.K., Chaturvedi, R., Drachenberg, C.B., Boucher, J.L., Hacker, A., et al. (2004). Protective role of arginase in a mouse model of colitis. J Immunol 173, 2109–2117. 10.4049/jimmunol.173.3.2109.

38. El Kasmi, K.C., Qualls, J.E., Pesce, J.T., Smith, A.M., Thompson, R.W., Henao-Tamayo, M., Basaraba, R.J., Konig, T., Schleicher, U., Koo, M.S., et al. (2008). Toll-like receptor-induced arginase 1 in macrophages thwarts effective immunity against intracellular pathogens. Nat Immunol 9, 1399–1406. 10.1038/ni.1671.

39. Sonoki, T., Nagasaki, A., Gotoh, T., Takiguchi, M., Takeya, M., Matsuzaki, H., and Mori, M. (1997). Coinduction of nitric-oxide synthase and arginase I in cultured rat peritoneal macrophages and rat tissues in vivo by lipopolysaccharide. J Biol Chem 272, 3689–3693. 10.1074/jbc.272.6.3689.

40. Haydar, D., Gonzalez, R., Garvy, B.A., Garneau-Tsodikova, S., Thamban Chandrika, N., Bocklage, T.J., and Feola, D.J. (2021). Myeloid arginase-1 controls excessive inflammation and modulates T cell responses in Pseudomonas aeruginosa pneumonia. Immunobiology 226, 152034. 10.1016/j.imbio.2020.152034.

41. Krischel, V., Bruch-Gerharz, D., Suschek, C., Kroncke, K.D., Ruzicka, T., and Kolb-Bachofen, V. (1998). Biphasic effect of exogenous nitric oxide on proliferation and differentiation in skin derived keratinocytes but not fibroblasts. J Invest Dermatol 111, 286–291. 10.1046/j.1523-1747.1998.00268.x.

42. Gallala, H., Macheleidt, O., Doering, T., Schreiner, V., and Sandhoff, K. (2004). Nitric oxide regulates synthesis of gene products involved in keratinocyte differentiation and ceramide metabolism. Eur J Cell Biol 83, 667–679. 10.1078/0171-9335-00425.

43. Isome, M., Lortie, M.J., Murakami, Y., Parisi, E., Matsufuji, S., and Satriano, J. (2007). The antiproliferative effects of agmatine correlate with the rate of cellular proliferation. Am J Physiol Cell Physiol 293, C705–711. 10.1152/ajpcell.00084.2007.

44. Nakamura, A., Kurihara, S., Takahashi, D., Ohashi, W., Nakamura, Y., Kimura, S., Onuki, M., Kume, A., Sasazawa, Y., Furusawa, Y., et al. (2021). Symbiotic polyamine metabolism regulates epithelial proliferation and macrophage differentiation in the colon. Nat Commun 12, 2105. 10.1038/s41467-021-22212-1.

45. Zahedi, K., Barone, S., and Soleimani, M. (2022). Polyamines and Their Metabolism: From the Maintenance of Physiological Homeostasis to the Mediation of Disease. Med Sci (Basel) 10. 10.3390/medsci10030038.

46. Hardman, M.J., and Ashcroft, G.S. (2008). Estrogen, not intrinsic aging, is the major regulator of delayed human wound healing in the elderly. Genome Biol 9, R80. 10.1186/gb-2008-9-5-r80.

47. Williams, H., Campbell, L., Crompton, R.A., Singh, G., McHugh, B.J., Davidson, D.J., McBain, A.J., Cruickshank, S.M., and Hardman, M.J. (2018). Microbial Host Interactions and Impaired Wound Healing in Mice and Humans: Defining a Role for BD14 and NOD2. J Invest Dermatol 138, 2264–2274. 10.1016/j.jid.2018.04.014.

48. Norman, G., Shi, C., Westby, M.J., Price, B.L., McBain, A.J., Dumville, J.C., and Cullum, N. (2021). Bacteria and bioburden and healing in complex wounds: A prognostic systematic review. Wound Repair Regen 29, 466–477. 10.1111/wrr.12898.

49. Williams, H., Crompton, R.A., Thomason, H.A., Campbell, L., Singh, G., McBain, A.J., Cruickshank, S.M., and Hardman, M.J. (2017). Cutaneous Nod2 Expression Regulates the Skin Microbiome and Wound Healing in a Murine Model. J Invest Dermatol 137, 2427–2436. 10.1016/j.jid.2017.05.029.

50. Anderson, C.J., Medina, C.B., Barron, B.J., Karvelyte, L., Aaes, T.L., Lambertz, I., Perry, J.S.A., Mehrotra, P., Goncalves, A., Lemeire, K., et al. (2021). Microbes exploit death-induced nutrient release by gut epithelial cells. Nature 596, 262–267. 10.1038/s41586-021-03785-9.

51. Andrabi, S.M., Sharma, N.S., Karan, A., Shahriar, S.M.S., Cordon, B., Ma, B., and Xie, J. (2023). Nitric Oxide: Physiological Functions, Delivery, and Biomedical Applications. Adv Sci (Weinh) 10, e2303259. 10.1002/advs.202303259.

52. Hall, J.R., Rouillard, K.R., Suchyta, D.J., Brown, M.D., Ahonen, M.J.R., and Schoenfisc, M.H. (2020). Mode of nitric oxide delivery affects antibacterial action. ACS Biomater Sci Eng 6, 433–441. 10.1021/acsbiomaterials.9b01384.

53. Estes Bright, L.M., Chug, M.K., Thompson, S., Brooks, M., Brisbois, E.J., and Handa, H. (2024). Analysis of the broad-spectrum potential of nitric oxide for antibacterial activity against clinically isolated drug-resistant bacteria. J Biomed Mater Res B Appl Biomater 112, e35442. 10.1002/jbm.b.35442.

54. Bila, I., Dzydzan, O., Brodyak, I., and Sybirna, N. (2019). Agmatine Prevents Oxidative-nitrative Stress in Blood Leukocytes Under Streptozotocin-induced Diabetes Mellitus. Open Life Sci 14, 299–310. 10.1515/biol-2019-0033.

55. Feng, Y., Piletz, J.E., and Leblanc, M.H. (2002). Agmatine suppresses nitric oxide production and attenuates hypoxic-ischemic brain injury in neonatal rats. Pediatr Res 52, 606–611. 10.1203/00006450-200210000-00023.

56. Zhu, H.E., Yin, J.Y., Chen, D.X., He, S., and Chen, H. (2019). Agmatinase promotes the lung adenocarcinoma tumorigenesis by activating the NO-MAPKs-PI3K/Akt pathway. Cell Death Dis 10, 854. 10.1038/s41419-019-2082-3.

57. Bachrach, U., and Weinstein, A. (1970). Effect of aliphatic polyamines on growth and macromolecular syntheses in bacteria. J Gen Microbiol 60, 159–165. 10.1099/00221287-60-2-159.

58. Liu, Z., Hossain, S.S., Morales Moreira, Z., and Haney, C.H. (2022). Putrescine and Its Metabolic Precursor Arginine Promote Biofilm and c-di-GMP Synthesis in Pseudomonas aeruginosa. J Bacteriol 204, e0029721. 10.1128/JB.00297-21.

59. Stover, C.K., Pham, X.Q., Erwin, A.L., Mizoguchi, S.D., Warrener, P., Hickey, M.J., Brinkman, F.S., Hufnagle, W.O., Kowalik, D.J., Lagrou, M., et al. (2000). Complete genome sequence of Pseudomonas aeruginosa PAO1, an opportunistic pathogen. Nature 406, 959–964. 10.1038/35023079.

60. Shah, P., and Swiatlo, E. (2008). A multifaceted role for polyamines in bacterial pathogens. Mol Microbiol 68, 4–16. 10.1111/j.1365-2958.2008.06126.x.

61. Williams, B.J., Du, R.H., Calcutt, M.W., Abdolrasulnia, R., Christman, B.W., and Blackwell, T.S. (2010). Discovery of an operon that participates in agmatine metabolism and regulates biofilm formation in Pseudomonas aeruginosa. Mol Microbiol 76, 104–119. 10.1111/j.1365-2958.2010.07083.x.

62. Tao, Y., Zheng, D., Zou, W., Guo, T., Liao, G., and Zhou, W. (2024). Targeting the cysteine biosynthesis pathway in microorganisms: Mechanism, structure, and drug discovery. Eur J Med Chem 271, 116461. 10.1016/j.ejmech.2024.116461.

63. Tikhomirova, A., Rahman, M.M., Kidd, S.P., Ferrero, R.L., and Roujeinikova, A. (2024). Cysteine and resistance to oxidative stress: implications for virulence and antibiotic resistance. Trends Microbiol 32, 93–104. 10.1016/j.tim.2023.06.010.

64. Szondi, D.C., Crompton, R.A., Oon, L., Subramaniam, N., Sze Han, L., Williams, H., Pennock, J., Lim, T.C., Wong, J., Vardy, L.A., and Cruickshank, S.M. (2024). A skin barrier role for Arginase in epithelial differentiation. [Manuscript in review].

65. Takahashi, M., Umehara, Y., Yue, H., Trujillo-Paez, J.V., Peng, G., Nguyen, H.L.T., Ikutama, R., Okumura, K., Ogawa, H., Ikeda, S., and Niyonsaba, F. (2021). The Antimicrobial Peptide Human beta-Defensin-3 Accelerates Wound Healing by Promoting Angiogenesis, Cell Migration, and Proliferation Through the FGFR/JAK2/STAT3 Signaling Pathway. Front Immunol 12, 712781. 10.3389/fimmu.2021.712781.

66. Niyonsaba, F., Ushio, H., Nakano, N., Ng, W., Sayama, K., Hashimoto, K., Nagaoka, I., Okumura, K., and Ogawa, H. (2007). Antimicrobial peptides human beta-defensins stimulate epidermal keratinocyte migration, proliferation and production of proinflammatory cytokines and chemokines. J Invest Dermatol 127, 594–604. 10.1038/sj.jid.5700599.

67. Ashcroft, G.S., Mills, S.J., Lei, K., Gibbons, L., Jeong, M.J., Taniguchi, M., Burow, M., Horan, M.A., Wahl, S.M., and Nakayama, T. (2003). Estrogen modulates cutaneous wound healing by downregulating macrophage migration inhibitory factor. J Clin Invest 111, 1309–1318. 10.1172/JCI16288.

68. Emmerson, E., Campbell, L., Davies, F.C., Ross, N.L., Ashcroft, G.S., Krust, A., Chambon, P., and Hardman, M.J. (2012). Insulin-like growth factor-1 promotes wound healing in estrogen-deprived mice: new insights into cutaneous IGF-1R/ERalpha cross talk. J Invest Dermatol 132, 2838–2848. 10.1038/jid.2012.228.

69. Dickson, M.A., Hahn, W.C., Ino, Y., Ronfard, V., Wu, J.Y., Weinberg, R.A., Louis, D.N., Li, F.P., and Rheinwald, J.G. (2000). Human keratinocytes that express hTERT and also bypass a p16(INK4a)-enforced mechanism that limits life span become immortal yet retain normal growth and differentiation characteristics. Mol Cell Biol 20, 1436–1447.

70. Rheinwald, J.G., Hahn, W.C., Ramsey, M.R., Wu, J.Y., Guo, Z., Tsao, H., De Luca, M., Catricalà, C., and O’Toole, K.M. (2002). A two-stage, p16(INK4A)- and p53-dependent keratinocyte senescence mechanism that limits replicative potential independent of telomere status. Mol Cell Biol 22, 5157–5172.

